# SV40 polyomavirus activates the Ras-MAPK signaling pathway for vacuolization, cell death, and virus release

**DOI:** 10.1101/454850

**Authors:** Nasim Motamedi, Xaver Sewald, Yong Luo, Walther Mothes, Daniel DiMaio

## Abstract

Polyomaviruses are a family of small, non-enveloped DNA viruses that can cause severe disease in immunosuppressed individuals. Studies with SV40, a well-studied model polyomavirus, have revealed the role of host proteins in polyomavirus entry and trafficking to the nucleus, viral transcription and DNA replication, and cell transformation. In contrast, little is known about host factors or cellular signaling pathways involved in the late steps of productive infection leading to polyomavirus release. We previously showed that cytoplasmic vacuolization, a characteristic late cytopathic effect of SV40, depends on the specific interaction between the major viral capsid protein VP1 and its cell surface ganglioside receptor GM1. Here we show that late during infection, SV40 activates a signaling cascade in permissive CV-1 monkey cells involving Ras, Rac1, MKK4 and JNK to induce SV40-specific cytoplasmic vacuolization and subsequent cell lysis and virus release. Inhibition of individual components of this signaling pathway inhibits vacuolization, lysis and virus release, even though high-level intracellular virus replication occurs. The identification of this pathway for SV40-induced vacuolization and virus release provides new insights into the late steps of non-enveloped virus infection and reveals potential drug targets for the treatment of diseases caused by these viruses.

**IMPORTANCE:** The polyomaviruses are small DNA viruses that include important model viruses and human pathogens that can cause fatal disease, including cancer, in immunosuppressed individuals. There are no vaccines or specific antiviral agents for any polyomavirus. Here, we show that late during infection, SV40 activates a signaling cascade involving Ras, Rac, and JNK that is required for cytoplasmic vacuolization and efficient virus release. This pathway may represent a new point of intervention to control infection by these viruses.

## INTRODUCTION

The polyomaviruses are small, non-enveloped, double-stranded DNA tumor viruses that include pathogenic human viruses such as BK polyomavirus (BKPyV), JC polyomavirus (JCPyV), and Merkel Cell polyomavirus (MCPyV), as well as the extensively studied model viruses, murine polyomavirus and the simian virus, SV40. MCPyV, the most recently discovered human tumor virus, is responsible for most cases of Merkel cell carcinoma, a rare but aggressive form of skin cancer. BKPyV is associated with inflammation of the urogenital tract and nephropathy, which can result in organ loss in renal transplant patients, as well as hemorrhagic cystitis in bone marrow transplant recipients [reviewed in (1)]. JCPyV is the causative agent of progressive multifocal leukencephalopathy (PML), a rare but usually fatal central nervous system demyelinating disease in immunocompromised individuals or patients receiving immunomodulatory monoclonal antibody treatment for various disorders (2). JCPyV and BKPyV infections are common in the human population. These viruses are phylogenetically closely related to SV40, which can cause PML-like brain pathology in immunosuppressed monkeys (3, 4). Therefore, SV40 serves as a model to study human polyomavirus pathogenesis, including neurological disease.

Productive polyomavirus infection of permissive cells can be divided into early and late phases. The early steps include virus binding to the cell surface, entry of virus particles into the cell, and trafficking of the viral genome to the nucleus where viral gene expression and DNA replication occur. Polyomavirus entry is initiated by binding of the major capsid protein VP1 to carbohydrate motifs on cell surface molecules. In the case of SV40, the ganglioside GM1 serves as the cellular receptor for infection. After endocytosis, polyomaviruses are transported to the endoplasmic reticulum (ER), where host factors initiate the disassembly of capsids and translocation of the viral genome and residual capsid into the cytoplasm for transport into the nucleus (5, 6). Expression of the early viral proteins including Large and small T antigen is followed by viral DNA replication, expression of the late viral proteins including VP1, and capid assembly, which occurs primarily in the nucleus before cells are lysed and mature infectious virus particles are released.

The initial interaction of a variety of polyomaviruses with cells acutely induces transient cellular signaling that supports the early steps of infection. JC virus induces ERK phosphorylation within minutes after receptor binding (7), which is required for the early stages of infection (8). Within the first two hours of infection, murine polyomavirus induces phosphoinositide 3’ kinase and Fak signaling pathways through binding of VP1 to gangliosides and α4-integrin (9-11). Inhibition of these signaling events can inhibit the early steps of murine polyomavirus infection. This virus also induces a second, delayed wave of mitogenic signaling that depends on viral early gene expression (11). Cell signaling also modulates productive SV40 infection (12). Binding of SV40 to GM1 at the plasma membrane triggers activation of more than 50 different kinases regulating the early steps of SV40 infection including local activation of tyrosine kinases to reorganize actin filaments for caveolin-1- or lipid raft-dependent SV40 internalization (13).

In contrast to the early stages of polyomavirus infection, late events leading to the release of virus particles are poorly understood. Viral proteins have been reported to facilitate SV40 release from cells. The late protein VP4 was reported to function as a viroporin with membrane-destabilizing properties that facilitates virus release, but these results have recently been challenged (14, 15). Furthermore, since VP4 is mostly found within the nucleus of infected cells, the mechanism leading to plasma membrane perforation and virus release is unclear. The minor capsid proteins VP2 and VP3 were also shown to support membrane permeabilization for virus release (14). These proteins can insert into or disrupt membranes when ectopically over-expressed in prokaryotic as well as eukaryotic cells.

SV40 infection of African green monkey cells leads to the appearance of characteristic cytoplasmic vacuoles late during infection, a phenomenon that led to the discovery of this virus in 1960 (16). We recently showed that vacuolization is triggered by the interaction between VP1 and GM1 at the cell surface (17, 18). SV40-induced vacuolization typically occurs late in infection. However, if large amounts of SV40 are added to cells, vacuoles can form acutely (17, 19). Virus replication is not required for vacuole formation, and purified VP1 pentamers are sufficient to induce vacuole formation. We hypothesized that the VP1-GM1 interaction triggers an as-yet-unidentified signaling cascade resulting in vacuolization (17).

Extensive cell vacuolization has also been observed in other experimental systems. Pore-forming toxins of various pathogens can induce the formation of cellular vacuoles and cell death (20, 21). Different types of intrinsic cell death programs, such as paraptosis and methuosis, are also associated with vacuole formation (20, 22). Cell signaling pathways including the Ras-MAPK pathway have been shown to contribute to vacuolization and non-apoptotic cell lysis in these processes (22, 23). However, cellular factors or signaling pathways have not been identified that are involved in vacuolization or other late events during SV40 infection.

In this study, we investigate the mechanism by which SV40 infection results in efficient virus release. We show that activation of the Ras-Rac1-MKK4-JNK signaling pathway late during SV40 infection results in vacuolization and ultimately facilitates cell lysis and release of progeny virus. Understanding the mechanism of polyomavirus release may allow the identification of proteins and pathways that can potentially be exploited as specific anti-viral drug targets for polyomaviruses and possibly other pathogenic, non-enveloped viruses.

## RESULTS

### Phenotypic characterization of SV40-induced vacuoles

We recently demonstrated that SV40-induced vacuole formation is triggered by binding of oligomeric VP1 to GM1 (17). Vacuolization typically occurs late during infection, but a detailed analysis of vacuole formation and its consequences for SV40 infection is lacking. We conducted physiological experiments, immune staining and live cell imaging to better characterize SV40-induced vacuoles. Vacuoles present 48 h.p.i. displayed an endocytic character as demonstrated by the rapid uptake of fluorescent low-molecular weight dextran (3kDa Dextran-Alexa488) from the culture medium (Fig. 1A). The endosomal nature of vacuoles was supported by immunostaining infected cells with antibodies against the early and late endosomal proteins EEA1 and Rab7, respectively, which revealed staining distributed around the circumference of vacuoles (Fig. 1B), presumably indicating the presence of these proteins of the vacuolar membrane. Although strong aggregation of the endoplasmic reticulum (ER) protein BiP was observed after infection, indicative of a cellular stress response, BiP was absent from vacuoles, suggesting that SV40-induced vacuoles are not composed of ER membranes (Fig. 1B).

**Figure 1.**
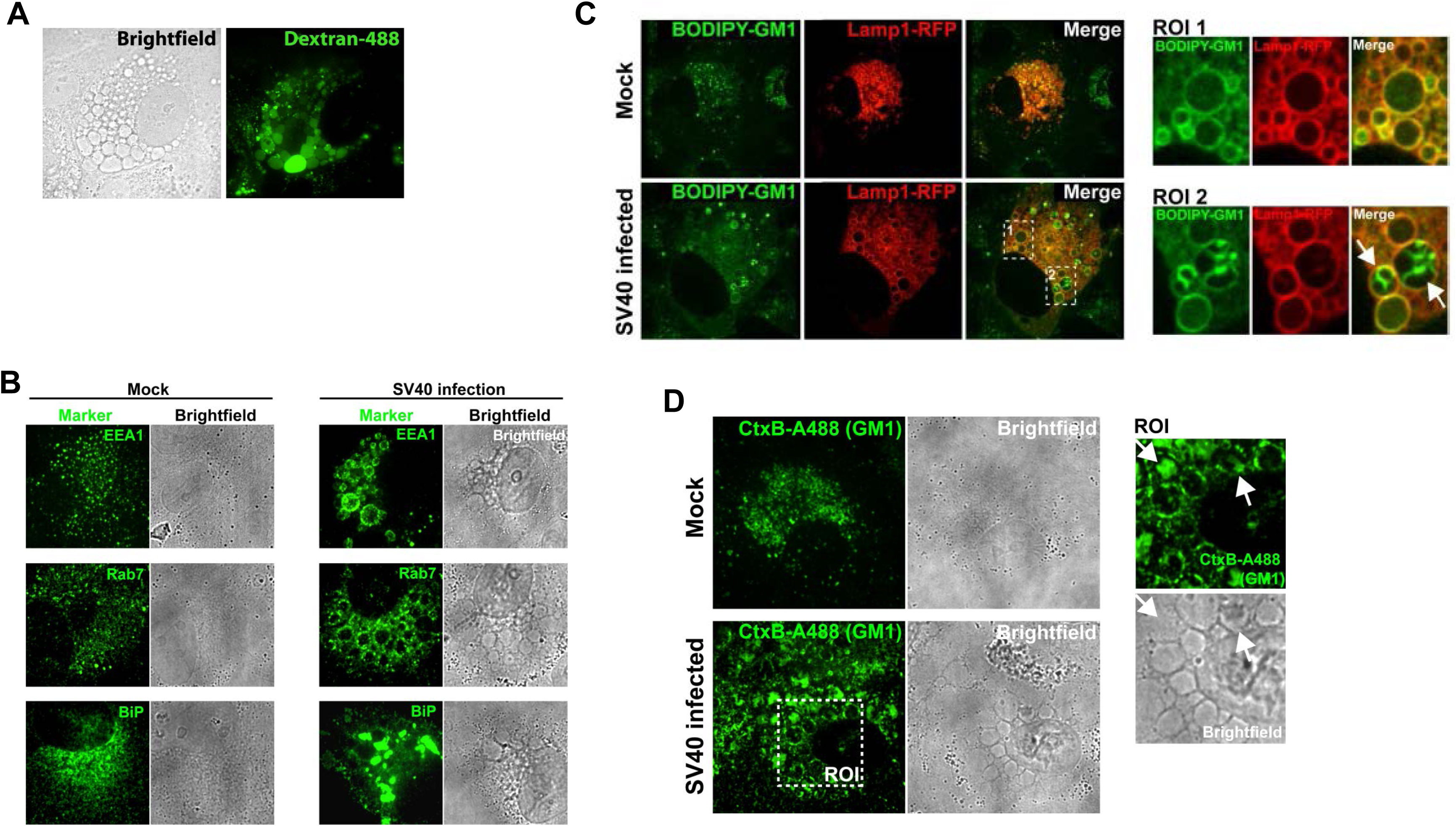
Characterization of SV40-induced vacuoles. (**A**) Corresponding representative fluorescence and bright-field images of SV40-infected CV-1 cells 48 h.p.i. after incubation with medium containing fluorescent Dextran-A488 (green). (**B**) Immunostaining and brightfield images of mock-infected and SV40-infected CV-1 cells 48 h.p.i. with antibodies recognizing markers of early endosome (EEA1), late (Rab7) endosome, and the endoplasmic reticulum (BiP), as indicated. (**C**) Fluorescence microscopy images of mock-infected and SV40-infected CV-1 cells pulse-labeled 48 h.p.i. with fluorescent GM1 (BODIPY-GM1, green). CV-1 cells expressing Lamp1-RFP (red) were visualized by confocal microscopy. Regions of interest (ROIs) 1 and 2 highlight vacuoles in infected cells showing BODIPY-GM1 in the limiting membranes and interior of Lamp1-positive vacuoles, respectively. Single planes of z-stacks are shown. (**D**) Fluorescence confocal microscopy and brightfield images of endogenous GM1 stained with fluorescent cholera toxin B (CtxB-Alexa488) (green) in mock-infected and SV40-infected CV-1 cells 48 h.p.i. Single planes of z-stacks are shown. ROI depicts CtxB-staining of vacuole membranes and intravacuolar GM1 in infected cells.

Because vacuole formation is triggered by the interaction between VP1 and GM1, we assessed whether GM1 is present in vacuoles. To determine the localization of GM1 during SV40-induced vacuole formation late during infection, infected and uninfected CV-1 cells expressing fluorescently-tagged Lamp1-RFP fusion protein were treated with fluorescently-labelled GM1 (BODIPY-GM1) and analysed by confocal microscopy. In contrast to mock-infected cells where GM1 displayed diffuse punctate staining, in infected cells GM1 (as well as the late endosomal/lysosomal marker Lamp1) was present at the limiting membrane of vacuoles (Fig. 1C). Endogenous GM1 displayed a similar distribution in vacuolar membranes 48 h.p.i. as assessed by staining with fluorescent cholera toxin B (CtxB-Alexa488) which, like VP1, binds to GM1 (Fig. 1D). In addition, strong patches of GM1 were present within some vacuoles (arrows) (Figs. 1C and 1D, regions of interest (ROI)), suggesting the existence of complex vesicular structures containing GM1 in infected cells.

To visualize the dynamics of vacuole formation and maturation, we conducted live-cell imaging using spinning-disk confocal microscopy of SV40-infected CV-1 cells transiently co-expressing fluorescently-tagged versions of the early endosome marker YFP-Rab5 and the lysosome marker Lamp1-RFP. Both marker proteins localized to vacuole membranes (Fig. 2). Fusion of Rab5-positive vacuoles was observed, indicating that the formation of large vacuoles was the result of fusion, not osmotic vesicle swelling (Fig. 2A; Movie 1). In addition, some Rab5-positive vacuoles matured into vacuoles containing Lamp1 (Fig. 2B; Movie 2), indicating that a dynamic endosomal system was involved in vacuole formation.

**Figure 2.**
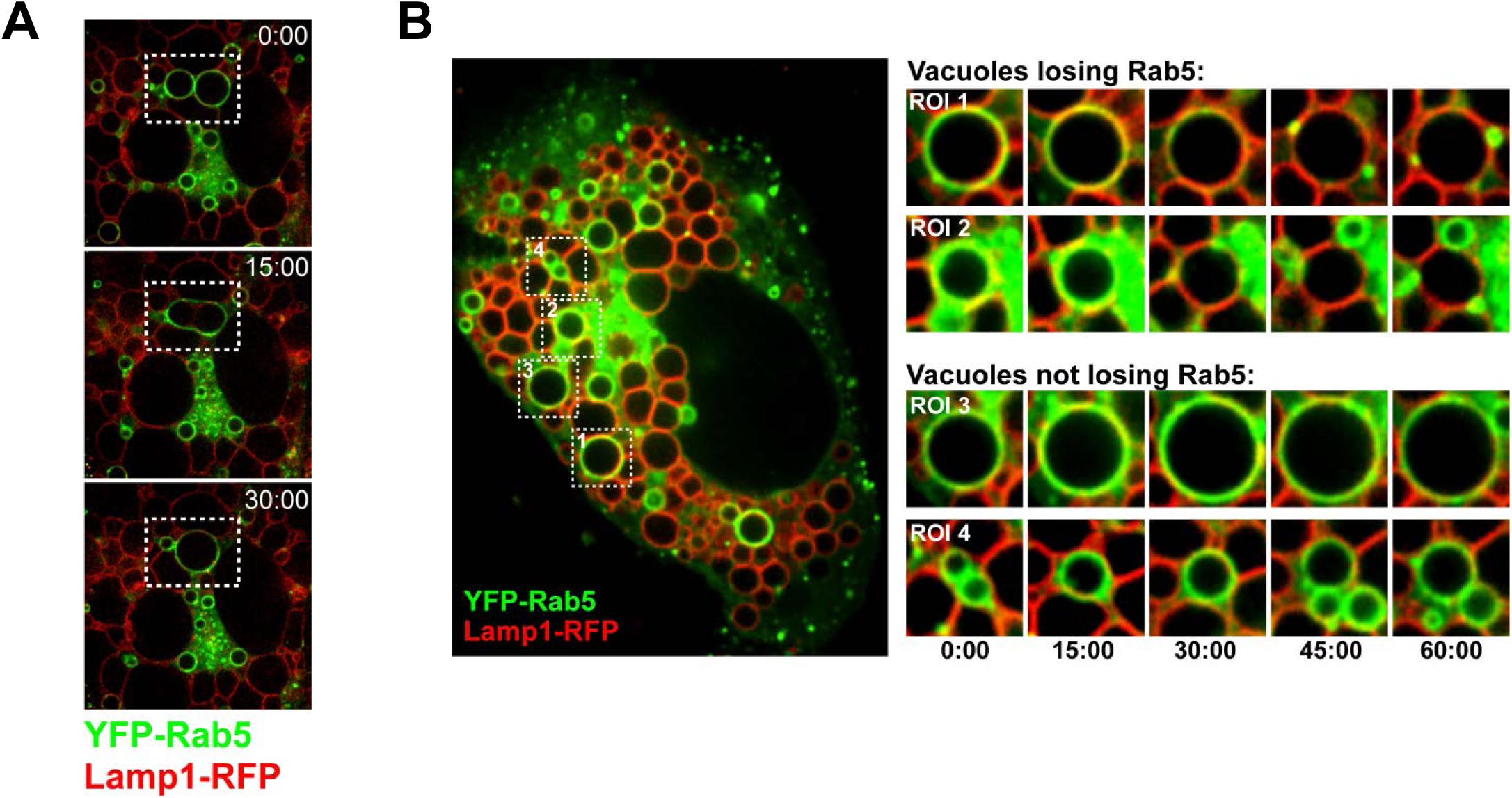
Dynamic vacuole formation. (**A**) Image sequence (top to bottom) from a time-lapse movie (Movie S1) showing fusion of YFP-Rab5-positive vacuoles in an SV40-infected CV-1 cell. Lamp1-RFP is shown in red. Boxes outline two YFP-Rab5 vacuoles that fuse. Numbers in this and panel B show time in minutes. (**B**) Overview image and time series of four regions of interest (ROI 1 to 4) from time-lapse movie S2 showing YFP-Rab5 (green) dynamics on SV40-induced vacuoles (Movie S2). ROIs 1 and 2 show vacuoles that lose YFP-Rab5 fluorescence and ROIs 3 and 4 show vacuoles with stable YFP-Rab5 fluorescence. Lamp1-RFP is shown in red.

### Vacuole formation requires Ras activity

Expression of activated Ras can lead to vacuole formation in glioblastoma and other cancer cell lines (22). To test for a role of Ras in SV40-induced cell vacuolization, we used a dominant-negative (DN) form of Harvey-Ras (HRas S17N), which inhibits the activation of all three Ras isoforms (H-, K- and N-Ras) (24). Plasmids expressing wild-type (WT) or DN versions of H-Ras, both fused to mEGFP, were transfected into CV-1 cells, which were infected 12 hours later with SV40. At 48 h p.i., Wild-type mEGFP-HRas co-localized to the membranes of vacuoles with VP1, but did not affect vacuolization as assessed by VP1 immunostaining and fluorescence microscopy (Fig. 3, upper panels and ROI). In contrast, expression of DN mEGFP-HRas S17N potently blocked SV40-induced vacuolization, even though VP1 was abundantly expressed (Fig. 3, lower panels). Flow cytometry of large T antigen expression in CV-1 cells gated for Ras-GFP expression revealed no difference in SV40 infection efficiency in cells expressing wild-type compared to DN mEGFP-HRas (Supplemental Fig. S1). These results indicate that Ras signalling is required for SV40-induced vacuole formation but not for SV40 infection.

**Figure 3.**
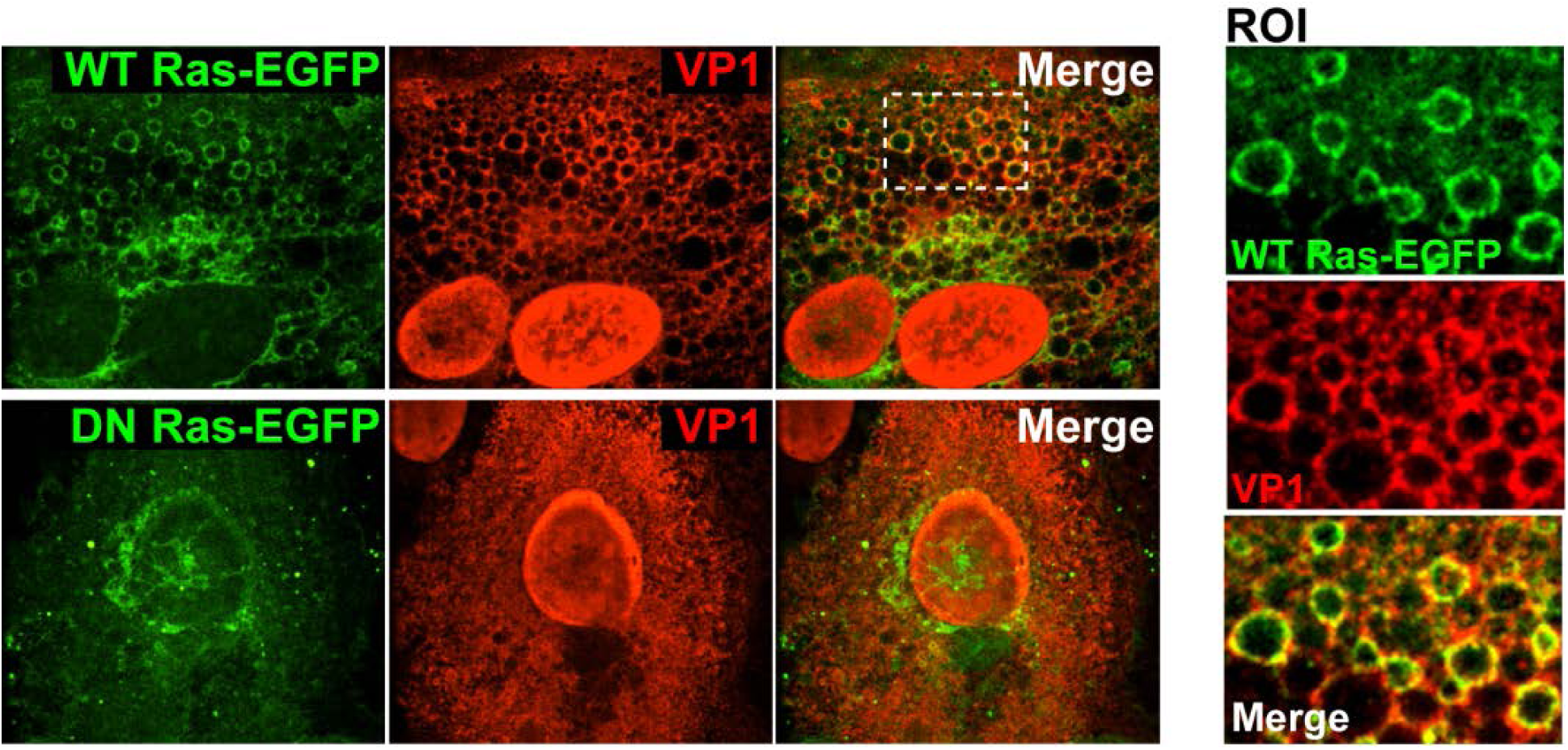
Ras signaling is required for vacuole formation. Fluorescence confocal microscopy images of SV40-infected CV-1 cells expressing wild-type (WT) or dominant-negative (DN) mEGFP-HRas (green). Forty-eight h.p.i., the localization of SV40 VP1 (red) was determined by immunostaining. The ROI depicts WT mEGFP-HRas accumulation at VP1-positive vacuoles. Single planes of z-stacks are shown.

### Vacuolization precedes cell lysis and SV40 release

To study the temporal relationship between SV40-induced vacuolization and progression of the virus life cycle, CV-1 cells were infected with wild-type SV40 at MOI 10, and vacuolization was monitored every 12 hours by bright-field microscopy. Small vacuoles first appeared at 36 h.p.i., with a pronounced vacuolization evident by 48 h.p.i. (Fig. 4A, C). Vacuolization reached a plateau at around 60 h.p.i. As expected, the appearance of vacuoles correlated with the expression of VP1, which became prominent around 36 h.p.i. as assessed by immunoblotting and grew stronger at later times (Fig. 4B).

**Figure 4.**
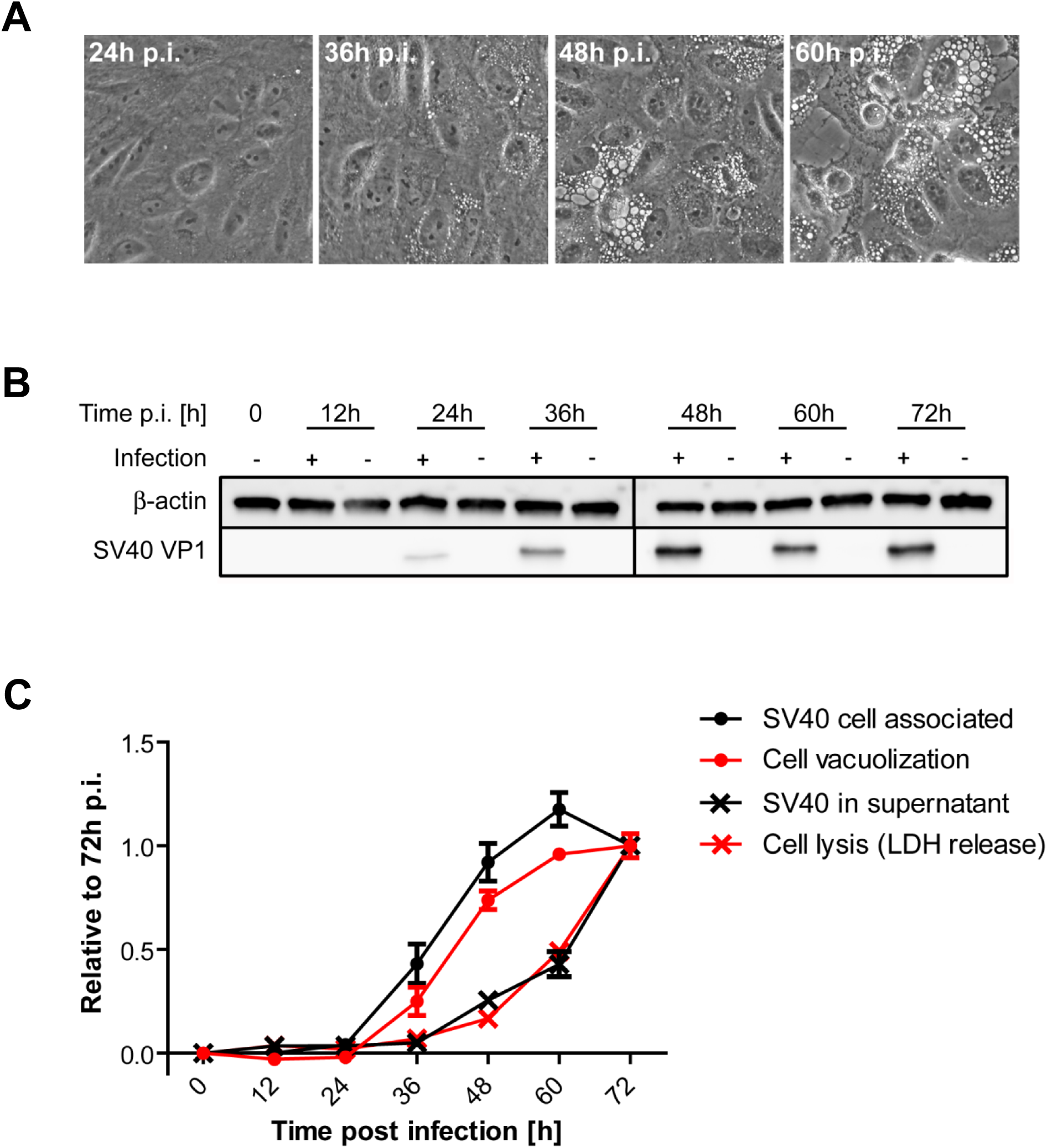
SV40-induced vacuole formation precedes cell lysis and virus release. (**A**) Time series of vacuole formation after infection of CV-1 cells with wild-type SV40 at MOI of 10. Vacuolization was monitored by bright-field microscopy at the indicated h.p.i. (**B**) Western blot analysis of VP1 and actin expression in mock- and SV40-infected CV-1 cells at the indicated times p.i. Mock-infected cells at each time point were used as control. (**C**) Quantitation of vacuolization, cell-associated SV40, cell lysis, and SV40 release over the time course of an SV40 infection. CV-1 cells were infected and the number of vacuolated cells at different time points after infection was quantified from bright-field images as depicted in **(A)**. A minimum of 200 cells per sample and three independent experiments were analyzed. Relative infectious units of cell-associated SV40 and released SV40 were quantified from cell lysates and supernatant, respectively, by titration onto CV-1 cells and flow cytometry analysis of large T antigen. CV-1 cell lysis was determined by quantitation of lactate dehydrogenase (LDH) in the supernatant using a colorimetric enzymatic assay, in which differences in the optical density between SV40-infected cells and mock-controls were determined. All values are displayed relative to 72 h.p.i. Mean +/- SEM from three independent experiments are shown.

To examine the temporal relationship between virus production and vacuole formation, we measured cell-associated and released infectious SV40. We infected CV-1 cells with SV40 at MOI of 10, and at various times p.i. the supernatant was collected as a source of released virus. At the same time points, the cells were lysed by freeze-thawing as a source of cell-associated virus. Infectious virus in both samples was quantified by infecting naïve CV-1 cells and enumerating large T antigen-positive cells by flow cytometry 24 h.p.i. As shown in Fig. 4C, the timing and extent of vacuolization was virtually congruent with the production of cell-associated virus, which first appeared at 36 h.p.i. In contrast, significant amounts of infectious SV40 in the supernatant were first detected 48 h.p.i. and continuously increased until the end of the observation period at 72 h.p.i. Finally, we assessed cell lysis by measuring the release of the intracellular enzyme lactate dehydrogenase (LDH) into the supernatant. LDH levels in the supernatant coincided with released SV40 (Fig. 4C). Thus, cell lysis and release of significant amounts of virus lag approximately 12 hours behind intracellular virus production and vacuolization.

### SV40 induces MAPK signaling late during infection

Infection by polyomaviruses at high MOI transiently activates cellular mitogenic signaling pathways, including the MAP kinase pathways, which are triggered by Ras activation (9-12). To determine whether the MAP kinase pathway was activated at late times after SV40 infection, when vacuoles typically form, we used phospho-site specific antibodies and western blotting to examine phosphorylation of the signaling proteins JNK, p38, and ERK. At 48 h.p.i., pronounced phosphorylation of JNK (Thr183/Tyr185), p38 (Thr180/Tyr182) and ERK (Thr202/Tyr204) was observed (Fig. 5A), with little difference in the total amount of these proteins, indicating broad activation of these signaling pathways in response to SV40 infection. We also examined the time course of MAP kinase signaling by analysing a series of time points beginning at 12 h.p.i., long after the acute phase of signaling has terminated. This analysis revealed the presence of progressive phosphorylation of JNK, ERK and p38 beginning as early as 24 h p.i. (Fig. 5B). Activation of signaling pathways at cell membranes can lead to JNK1/2 phosphorylation through the action of MKK4, a membrane-proximal kinase. To determine the phosphorylation status of the MKK4 during SV40 infection, we conducted western blot analysis of MKK4 phosphorylation at serine 257 and threonine 261. This analysis revealed the presence of phospho-MKK4 by 48 h.p.i. (Fig. 5B). Although MKK4 phosphorylation was detected at later times than JNK phosphorylation, we believe that this is due to the lower sensitivity of the phospho-MKK4 antibody. After phosphorylation, activated MAP kinases translocate into the nucleus and regulate gene expression by phosphorylating transcription factors such as ATF-2 and c-Jun. Consistent with activation of MAP kinase signaling, ATF-2 and c-Jun were phosphorylated late during SV40 infection, when VP1 expression became abundant (Fig. 5B).

**Figure 5.**
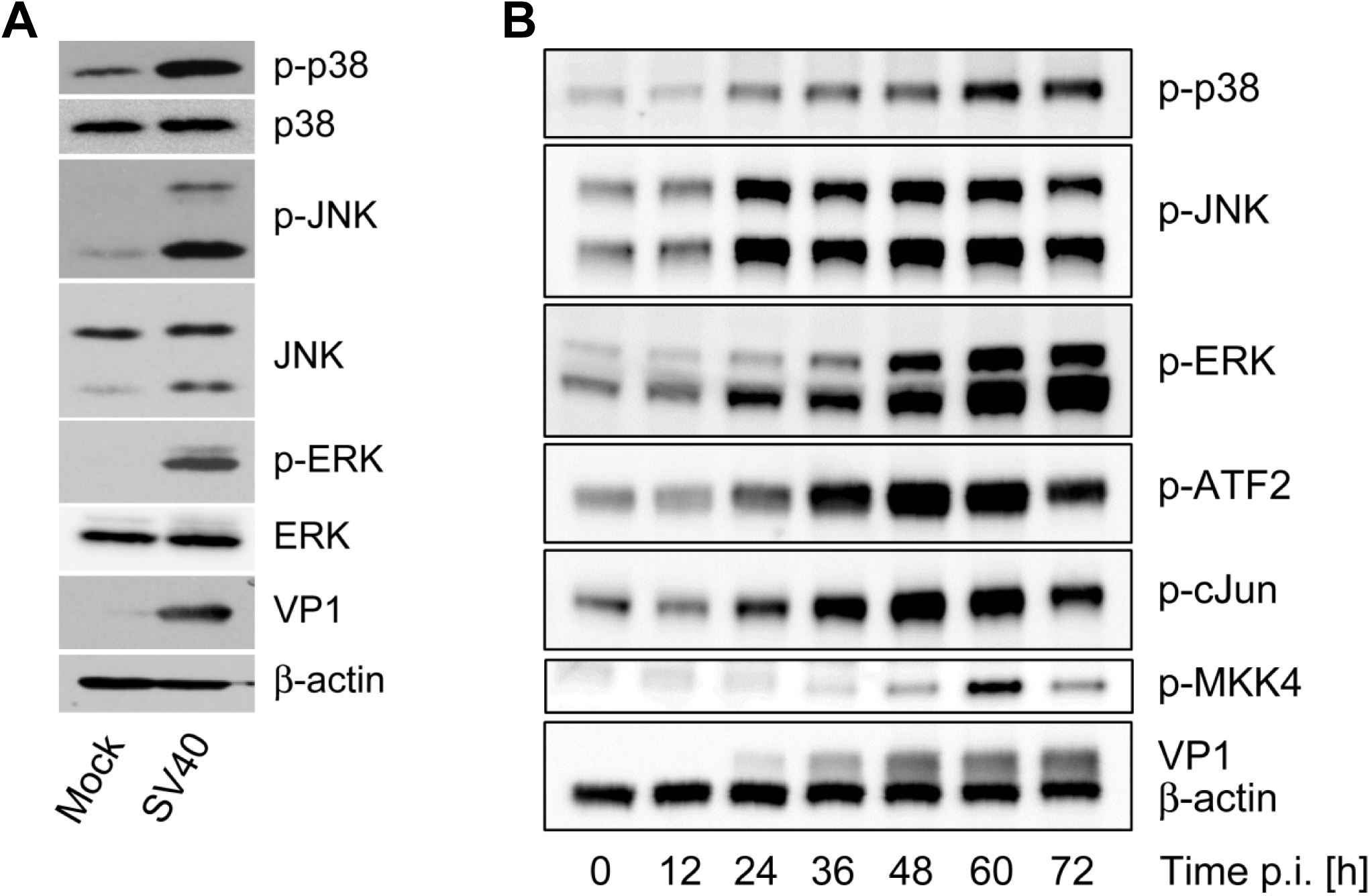
SV40 infection activates intracellular signaling pathways at late times after infection. (**A**) Western blot analysis of phosphorylated and total p38, JNK and ERK in mock-infected or SV40-infected CV-1 cells 48 h.p.i. VP1 and β-actin expression are shown as controls. (**B**) Western blot analysis of CV-1 cells over the time course of SV40 infection. Samples harvested at the indicated h.p.i. were analyzed for phosphorylated p38, JNK, ERK, MKK4, ATF2, and c-Jun, as well as for β-actin and VP1 expression.

### JNK, MKK4, and Rac1 are required for vacuolization and SV40 release

To determine the role of specific signaling pathways in vacuolization and SV40 release, we tested whether chemical and genetic inhibitors that blocked signaling affected these processes. Starting at 12 h.p.i., CV-1 cells were treated with chemical inhibitors targeting JNK (SP600125), p38 (SB203580) and MEK (Selumetinib), a key component of the ERK pathway. Inhibitory activity was confirmed by western blot analysis showing reduced target protein phosphorylation 48 h.p.i. (Fig. 6A). Vacuolization of cells treated with the inhibitors was examined two days p.i. Strikingly, the JNK inhibitor, SP600125, completely blocked vacuole formation, whereas p38 inhibition had no effect compared to vehicle alone (Figs. 6B and 6C). The ERK pathway inhibitor caused a significant reduction in vacuolization, but strong cytotoxic effects were observed during treatment of non-infected cells with this compound (Supplemental Fig. S2A). We conclude that the lack of vacuoles in ERK-inhibited cells is likely a consequence of accelerated cell death and detachment of cells rather than a true suppression of vacuolization.

**Figure 6.**
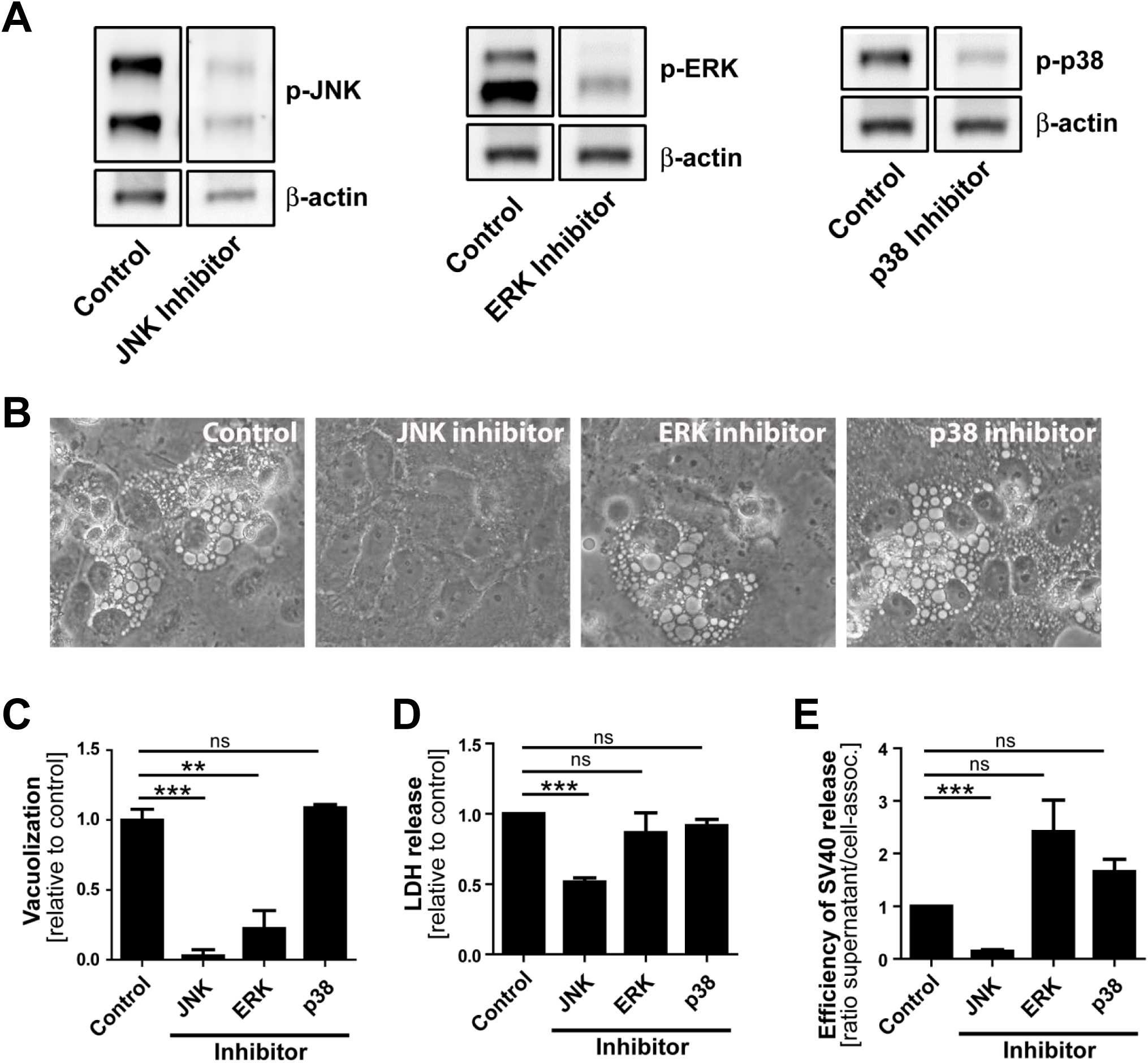
MAP kinase components are required for efficient vacuolization, cell lysis and virus release. (**A**) Inhibitors SP600125, Selumetinib, and SB203580 inhibit SV40-induced phosphorylation of JNK, ERK, and p38, respectively. CV-1 cells were infected at MOI of 10. Inhibitor treatment was started at 12 h.p.i. and immunoblotting was performed on extracts prepared 48 h.p.i. (**B**) Bright-field images of CV-1 cells 48 h.p.i. after infection with SV40. CV-1 cells were infected at MOI of 10 and treated with inhibitors against JNK, ERK, and p38 or DMSO vehicle at 12 h.p.i. (**C**) The number of vacuolated cells two days after infection was quantified and normalized to DMSO-treated control cells. (**D**) Analysis of cell lysis two days post-infection with SV40 in CV-1 cells treated with inhibitors. LDH activity in the supernatant was measured. **(E)** The ratio of released SV40 in supernatant versus cell-associated SV40 is shown. Data were normalized to DMSO-treated control cells. (Quantitation of cell-associated SV40 and SV40 in the supernatant 48 h.p.i. of CV-1 cells treated with JNK, ERK, and p38 inhibitors is shown in Supplemental Fig. S2.) The mean values +/- SEM from three independent experiments are shown.

Inhibition of JNK also inhibited SV40-induced cell lysis by 50% (Fig. 6D) and caused a 6-fold reduction of SV40 release with respect to cell-associated virus (Figs. 6E, S2B, and S2C). In contrast, treatment of infected cells with the MEK or p38 inhibitor did not reduce cell lysis or SV40 release. Treatment of cells with inhibitors did not interfere with early steps of SV40 infection as assessed by flow cytometry for large T antigen expression (Supplemental Fig. S2D), suggesting that the JNK signaling pathway is specifically required late in infection for vacuolization, cell lysis, and efficient virus release.

To assess the role of MKK4 in SV40-induced vacuolization and virus release, we used three shRNAs with different *mkk4* target sequences to generate CV-1 cells with stable MKK4 knock-down (Fig. 7A). MKK4 knock-down by each of these shRNAs reduced SV40-induced cell vacuolization (Figs. 7B and 7C) and cell lysis (Fig. 7D) compared to scrambled shRNA control. Notably, MKK4 knock-down also caused a significant reduction of SV40 release from infected cells compared to cell-associated SV40 (Fig. 7E; Supplemental Figs. S3A and S3B). MKK4 knock-down did not affect the efficiency of SV40 infection as assessed by intracellular virus production (Supplemental Fig. S3A), and by flow cytometry and western blotting for large T antigen and VP1 expression (Supplemental Figs. S3C and S3D).

JNK and MKK4 signaling is required for methuosis, a Ras-dependent form of cell death characterized by extensive vacuolization (25). The small GTPase Rac1 has been identified as a mediator of methuosis upstream of MKK4 and JNK. Therefore, we tested the role of Rac1 in SV40-induced vacuolization, cell lysis, and virus release. Treatment of CV-1 cells 12 h.p.i. with the specific Rac1 inhibitor EHT1864 blocked downstream phosphorylation of MKK4 compared to vehicle-treated cells, confirming inhibition of the Rac1-MKK4-JNK signaling cascade (Fig. 8A). Rac1 inhibition also significantly reduced cell vacuolization (Figs. 8B and 8C), cell lysis (Fig. 8D), and release of infectious SV40 (Fig. 8E; Supplemental Figs. S4A and S4B). Production of intracellular virus and SV40 infectivity were not inhibited by EHT1864 (Supplemental Figs. S4A and S4C). Taken together, these results support a model in which a Ras-dependent cell signaling cascade involving Rac1-MKK4-JNK induces extensive cell vacuolization, followed by cell lysis and SV40 release.

**Figure 7.**
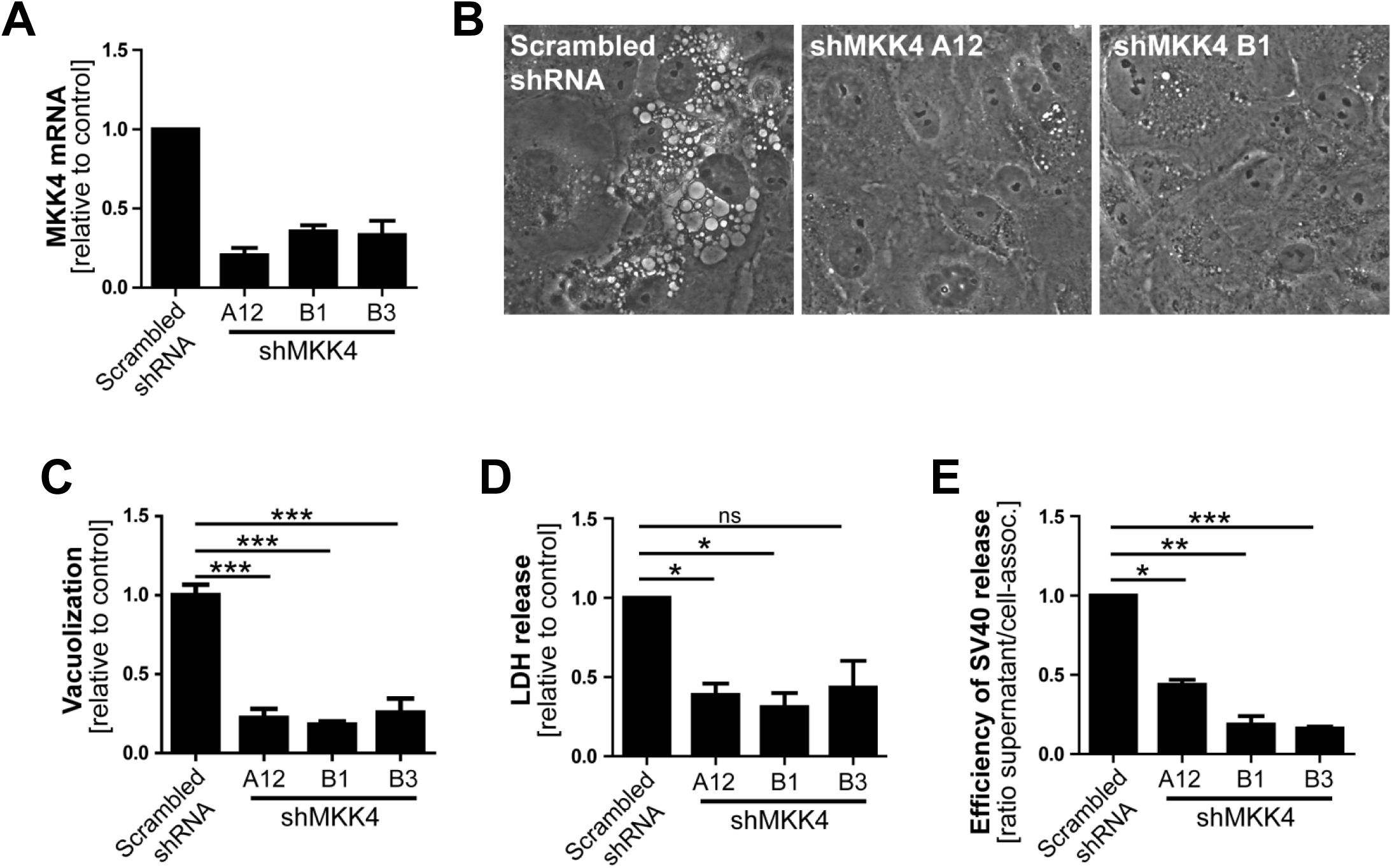
MKK4 is required for efficient SV40-induced vacuolization, cell lysis, and virus release. (**A**) qPCR analysis of *mkk4* mRNA expression levels in CV-1 cells stably expressing three different shRNAs targeting MKK4 (A12, B1, B3). Levels of mRNA were normalized to mRNA in control cells expressing scrambled shRNA. (**B**) Images of SV40-infected control and MKK4 knock-down cells. CV-1 cells stably expressing scrambled control shRNA or MKK4 A12 or B1 shRNA were infected with SV40, and vacuole formation was monitored by bright-field microscopy 48 h.p.i. Similar results were obtained with B3 shRNA. (**C**) The number of vacuolated cells as in panel B was quantified and normalized to scrambled shRNA control cells. A minimum of 200 cells per sample were analyzed. The mean values +/- SEM from three independent experiments are shown. (**D**) CV-1 cells expressing three different MKK4 shRNAs were infected with SV40 and the supernatants were analyzed for LDH release 48 h.p.i. **(E)** The ratio of released SV40 in supernatant versus cell-associated SV40 is shown. (Quantitation of cell-associated SV40 and SV40 in supernatant of MKK4 knock-down cells 48 h.p.i. is shown in Supplemental Figs. S3A and S3B.) Data were normalized to scrambled shRNA control cells.

**Figure 8.**
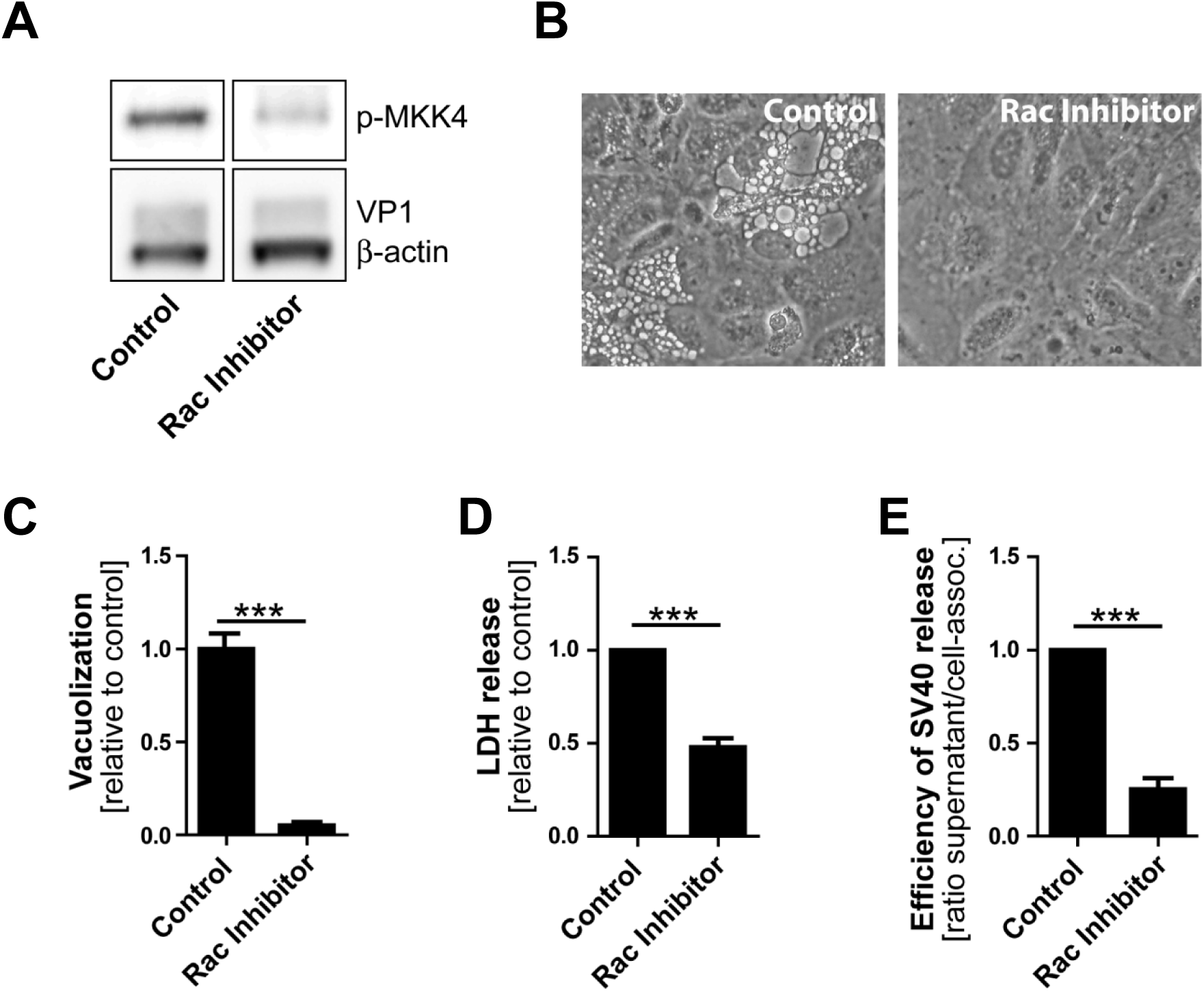
Rac1 activity is required for efficient SV40-induced vacuole formation, cell lysis, and virus release. CV-1 cells were infected with SV40 at MOI of 10. At 12 h.p.i., the Rac1 inhibitor EHT1864 was added and cells were analyzed at 48 h.p.i.(**A**) Western blot analysis of MKK4 phosphorylation in SV40-infected CV-1 cells in the presence and absence of Rac1 inhibitor. **(B)** Infected CV-1 cells were treated with EHT1864 or DMSO control and photographed by bright-field microscopy. (**C**) Quantitation of vacuolated CV-1 cells after SV40 infection as described in Fig. 6D. (**D**) Infected CV-1 cells were treated with EHT1864. 48 h.p.i. LDH released in the supernatant was determined. **(E)** The ratio of released SV40 in supernatant versus cell-associated SV40 is shown. To quantify virus release, infectious units of SV40 in cell lysates and supernatants were analyzed by infection and flow cytometry, as described in Figs. 2D-F. (Quantitation of cell-associated SV40 and SV40 in supernatant in Rac1 inhibitor-treated CV-1 cells 48 h.p.i. is shown in Supplemental Figs. S4A and S4B.)

## DISCUSSION

Activation of cellular signaling pathways is important during various steps in virus life cycles. For example, virus binding to its surface receptor often triggers signaling cascades that facilitate virus entry and replication. We previously reported that intracellular SV40 VP1 expression alone does not induce vacuolization and that SV40 particles and VP1 are absent from the vacuolar lumen in infected CV1 cells or CV1 cells undergoing vacuolization in response to acute treatment with VP1 pentamers (17). These findings suggest that vacuolization is the result of a signaling cascade triggered at the cell surface. In this report, we show that in addition to signaling occurring early in infection, SV40 also induces the MAPK signaling cascade during the late stage of infection when large amounts of VP1 accumulate to support vacuolization and efficient virus release.

Several findings reported here suggest that progeny SV40 particles bind to GM1 at the plasma membrane and trigger GM1-dependent Ras activation and vacuolization to support further virus release. Live cell microscopy of SV40-infected CV-1 cells revealed that SV40-induced vacuoles display a dynamic endocytic nature, consistent with vacuolization having a signaling basis. We also show that VP1 co-localizes with GM1 and Ras at the limiting membrane of SV40-induced vacuoles arising late in infection. Most importantly, we showed that dominant-negative Ras S17N or inhibition of JNK signaling inhibits vacuole formation. Overall, we hypothesize that in response to SV40-induced clustering of GM1, Ras is activated and triggers MAPK signaling through Rac1-MKK4-JNK, which results in vacuolization and subsequent cell lysis and virus release.

Previously published work is consistent with this model. GM1 clustering has been reported to modify active HRas distribution in membrane nanodomains (26), and overexpression of constitutively-active HRas (HRas G12V) leads to extensive vacuolization in glioblastoma cells, with Ras localized at the limiting membrane of cytoplasmic vacuoles (22, 23). Like vacuoles formed during SV40 infection, HRas-induced vacuole formation in glioblastoma cells was independent of ERK signaling (23) and blocked by chemical inhibition of Rac1(27). Furthermore, in both Ras-activated glioblastoma cells and SV40-infected cells, vacuoles contain markers such as Lamp-1 and undergo extracellular fluid uptake resembling macropinocytosis (22, 23, 28).

Although Ras and VP1 co-localize in vacuolar membranes (Fig. 1C), we have not determined the cellular compartment where GM1-dependent Ras activation occurs during SV40 infection. GM1 can localize to several different membrane compartments including the plasma membrane, Golgi network and the ER, and the ceramide structure of GM1 influences its trafficking into various cellular compartments (29). Similarly, Ras localizes not only to the plasma membrane but also to internal membranes such as the limiting membranes of endosomes, the ER and the Golgi network (30). FRET-based assays demonstrated that EGF stimulation changed the distribution of endogenous active Ras from the plasma membrane to endosomal-like intracellular vesicles, the Golgi network and the ER (31). Moreover, the subcellular localization of active Ras influences signaling through downstream effector pathways (32). Thus, SV40 may activate Ras in various cellular compartments at different steps during the viral life cycle, with different biological consequences.

Time course analysis of SV40-induced late cellular signaling revealed that it precedes vacuolization, cell lysis and virus release. Active phosphorylated forms of JNK were detected 24-36 h.p.i., coincident with VP1 expression, followed by phosphorylation of nuclear transcription factors. Unlike the acute transient signaling induced upon virus cell binding, this late phase signaling is sustained. Activation of cell signaling preceded vacuole formation with the first vacuoles emerging at 36-48 h.p.i., prior to cell lysis and virus release. Although other cellular kinases such as p38 and ERK were also activated during SV40 infection, inhibitor experiments revealed that only the JNK pathway is essential for vacuolization, cell lysis and efficient virus release. Importantly, inhibition of JNK, Ras, and MKK4 activity or expression did not interfere with SV40 entry and intracellular replication, so the impaired late events do not merely reflect an early replication block. This is consistent with an earlier published report that MAPK/ERK signaling is not required for the early response to SV40 infection (12) and with a recent genome-wide analysis of kinases that contribute to SV40 endocytosis, which did not detect a requirement for MAPK signaling (13). We conclude that Ras-Rac1-MKK4-JNK signaling is essential late during SV40 infection for vacuolization and cell death leading to virus release, whereas other signaling pathways are dispensable.

In agreement with previous studies (33), we showed that infectious SV40 first appears at low levels in the supernatant as early as 36 h.p.i. The release of virus at this early time might occur due to the first cells that lyse or to non-lytic virus release. We hypothesize that SV40 released from cells around this time binds to the plasma membrane of the same and neighboring infected cells and induces cell signaling, which in turn stimulates cell lysis and subsequent increased virus release from these cells, thus establishing a positive feedback loop that stimulates further signaling and virus release. This model is similar to our earlier analysis of vacuolization during SV40 infection (17), in which we proposed that the first released virus binds to cell surface GM1 and stimulates vacuole formation late during infection. We extend the model here to include virus release as well as vacuolization as a phenotype that can be acutely triggered late in infection by the first progeny virus released. Thus, the initial wave of released virus primes the infected cell population for more pronounced vacuolization and enhanced virus release.

To complete the virus life cycle, polyomaviruses release depends on lysis of the infected cell. Whether cell death is the consequence of plasma membrane rupture resulting from extensive virus production or a regulated process depending on active cellular signaling was heretofore unclear. Here, we show that high levels of intracellular virus are not sufficient for efficient release and that cellular MAPK signaling is necessary for optimal cell lysis and SV40 release. This lytic process resembles methuosis, which, as noted above, is a Rac1-dependent cell death pathway displaying characteristic cellular vacuolization occurring after ectopic expression of oncogenic Ras (22, 23, 34).

Vacuolization and virus release are both facilitated by MAPK signaling and vacuolization precedes virus release. If activation of the Ras-MAPK signaling cascade independently induces both vacuole formation and virus release, vacuolization is a convenient marker for the signaling events that foster efficient release. Alternatively, it is possible that signaling leads to vacuolization, and that vacuolization itself facilitates subsequent cell death and enhanced virus release.

In addition to the role of cellular signaling, virus encoded proteins could also be involved in cell lysis and SV40 release. VP4 is a late SV40 protein previously reported to function as a viroporin to support virus release which disrupted membranes when ectopically added to red blood cells, liposomes or Cos-7 cells (35). However, more recent studies did not confirm the lytic activity of VP4 during SV40 infection (15). In the context of SV40 replication, the expression of VP1 alone, without VP2 and VP3, leads to cell lysis and the release of viral particles, suggesting that lytic activity is mediated via VP1, likely through activation of a cellular program as reported here (14, 15).

Our results raise the possibility that JNK and MAPK pathway inhibitors may have a role in treating polyomavirus infections by decelerating virus propagation and spread within the host by reducing virus release. This would presumably provide a protective effect in affected tissues while immune reconstitution is underway (36). Because of the long replication cycle of human polyomaviruses and difficulties in synchronizing infection, late events are difficult to study. Nevertheless, further studies of the human pathogenic polyomaviruses may reveal that they are also affected by signaling programs late during infection. Although human polyomaviruses bind only weakly to GM1 (36), a variety of stimuli can activate the signaling elements described here. Thus, analysis of late events of human polyomavirus infection might establish that this or overlapping signaling pathways are viable therapeutic targets.

## MATERIAL and METHODS

### Cells and virus

CV-1 cells and SV40 776 virus DNA were purchased from American Type Culture Collection (ATCC). Cells were maintained in Dulbecco’s modified Eagle’s medium (DMEM) supplemented with 10% fetal bovine serum (FBS), 10 mM L-glutamine, and 10 mM HEPES (pH 7.2) in 5% CO_2_ at 37°C. SV40 was produced from SV40 776 in a bacterial vector backbone puc19 by excision, re-ligation and transfection into CV-1 cells. When significant cell death was observed, cell cultures were subjected to multiple rounds of freeze/thaw lysis. Cellular debris was removed by centrifugation at 1,000 rpm for 5 min, and supernatants were filtered through 0.45µm syringe filters, aliquoted, and stored at -80°C. To produce higher titer virus stocks, fresh CV-1 cells were infected at MOI of 0.5 and processed as described above.

### Virus titer quantitation

To quantify infectious units of SV40, serial dilutions of virus preparation, tissue culture supernatant or cell lysate were added to monolayers of 2 x 10^5^ CV-1 cells in six-well plates. After 24 h, CV-1 cells were trypsinized, fixed, and permeabilized in methanol or with 4% PFA/0.5% Triton X-100 before being subjected to immunofluorescence staining for large T antigen and flow cytometry. In a typical infection with wild-type SV40, at 48 h.p.i. there is approximately 5 to 9-fold more virus in the supernatant than in the cell lysate.

### Immunoblots

CV-1 cells were seeded at 2 x 10^5^ in six-well plates and infected with SV40 on the following day at MOI of 10. At indicated times p.i., cells were harvested by lysis in lysis buffer (2% Triton X-100, 0.5% Na-deoxycholate, 150 mM NaCl, 25 mM Tris, 5 mM EDTA, Halt protease and phosphatase inhibitors (Thermo Scientific)). Extracts were suspended in 5 x Laemmli buffer, subjected to sonication, and boiled. Equal sample volumes were loaded on SDS-PAGE for protein separation. Proteins were then transferred to 0.2 μm polyvinylidene difluoride (PVDF) membranes in Tris/glycine transfer buffer (25 mM Tris, 192 mM glycine, and 20% methanol) for 2 h at 100V. Membranes were blocked in 5% BSA/TBST or non-fat dry milk in TBST (10 mM Tris-HCl, pH 7.4, 167 mM NaCl, 1% Tween-20) for 1 h and incubated overnight at 4°C with indicated antibodies in 5% BSA/TBST for phosphorylated targets or 5% non-fat dry milk/TBST for all others. Blots were washed in TBST and incubated for 1 h at room temperature with HRP-conjugated donkey anti-mouse/rabbit/goat (Jackson ImmunoResearch) in 5% non-fat dry milk/TBST. After washing with TBST, blots were visualized by enhanced chemiluminescence (SuperSignal West Pico/Femto Chemiluminescent Substrate, Thermo Scientific [CST]). The following primary antibodies were used for immunoblotting: anti-ß-actin (Abcam, ab #8227); anti-p-ERK (Cell Signaling Technology, CST, #4370); anti-p-JNK (CST, #4668); anti-p-p38 (CST, #4511); anti-p-MKK4 (CST, #9156); anti-ERK (CST, #4695); anti-JNK (CST, #9252); anti-p38 (CST, #8690); anti-p-ATF2 (CST, #9225); anti-p-cJun (CST, #3270); anti-large T antigen (PAb, #108); anti-VP1 (PAb, #597).

### Vacuolization assay

CV-1 cells were plated at a density of 2 x 10^5^ cells per well on six-well plates and infected with SV40 on the following day at MOI of 10. At indicated time points, vacuolization was assessed and documented by bright-field microscopy. To quantify vacuolization, a minimum of 200 cells per sample were counted under blinded conditions.

### Chemical inhibitor treatment

CV-1 cells were plated at a density of 2 x 10^5^ on six-well plates. On the following day, cells were infected with SV40 at MOI of 10. Twelve h.p.i. or at the time of infection (0 h.p.i.), chemical inhibitors were added at a concentration of 25 µM to the infected and mock-treated CV-1 cells. Inhibitor treatment was maintained until the end of the experiment. The following chemical inhibitors were used in this study: Selumetinib (ERK pathway inhibitor, inhibits MEK1); EHT 1864 (Rac inhibitor); SB203580 (p38 inhibitor); SP600125 (JNK inhibitor). All inhibitors were purchased from Selleckchem.

### qRT-PCR

For quantitative PCR, total RNA was extracted from 2 x 10^5^ cells using the RNEasy Mini Kit (Qiagen) and a maximum of 1 µg cDNA was transcribed with the iScript™ cDNA Synthesis Kit (BioRad). The relative expression levels were assessed in triplicate on a single color detection system (BioRad CFX Connect Real-Time PCR Detection System) with the iTaq™ universal SYBR Green supermix (BioRad). Genes and primers used for qPCR were as follows: GAPDH FW (TGGTATCGTGGAAGGACTCA), GAPDH RV (CCAGTAGAGGCAGGGATGAT), MKK4 FW (TGAAAAGGCACAAAGTAAACGCA), MKK4 RV (CCCAGTGTTGTTCAGGGGAG).

### Dextran and BODIPY-GM1 treatment

CV-1 cells were plated at a density of 2 x 10^4^ on Lab-Tek II chambered coverglass slides (Nunc) and infected with SV40 at MOI of 100 on the following day. At 47 h.p.i., cells were incubated for 1 h with fluorescent markers. For dextran uptake assays, the cell culture medium was supplemented with 0.25 mg/ml of 3 kDa Dextran conjugated with Alexa Fluor 488 (Dextran-A488, Molecular Probes). For labeling with fluorescent GM1, 5 µM BODIPY FL C5-GM1 (Molecular Probes) was added to the cell culture medium. At 48 h.p.i., cells were thoroughly washed and fresh cell culture medium was added for imaging. Cells were imaged at 50 h.p.i. on a Nikon TE2000 spinning disk confocal microscope driven by the Volocity software package (Perkin Elmer).

### Immunofluorescence

CV-1 cells were plated at a density of 3 x 10^4^ on Millicell EZ Slide four-well glass slides (Millipore). On the following day, cells were infected with SV40 at MOI of 100. After 48 h, cells were fixed with 4% PFA, permeabilized with 0.5% Triton X-100 and immunostained or treated with 0.5 µg/ml Alexa Fluor 488-conjugated CtxB (Molecular Probes) to stain GM1. The following primary antibodies were used for immunofluorescence staining: anti-VP1 (Abcam, ab #53977); anti-EEA1 (CellSignaling, #C45B10); anti-Rab7 (CellSignaling, #D95F2); anti-BiP (Abcam, ab #108615). The secondary antibodies donkey anti-mouse/anti-rabbit conjugated to Alexa Fluor-488 or -568 (Molecular Probes) were used. Stained samples were embedded in ProLong Gold (Invitrogen) and data were acquired on a spinning disk confocal microscope (Nikon). Images were analysed using Volocity software (Perkin Elmer). For experiments involving Ras expression, 2 x 10^5^ CV-1 cells were plated on coverslips in six-well plates. Cells were transfected with WT or DN mEGFP-HRas using FuGENE6 (Promega) transfection reagent. On the following day, cells were infected with SV40 at MOI of 10. At 48 h.p.i., the cells were fixed and immunostained with a primary antibody against anti-VP1 (Abcam) and the secondary antibody anti-rabbit Alexa Fluor 568. Stained samples were embedded in ProLong Gold (Invitrogen) and data were acquired on a spinning disk confocal microscope (Nikon). Images were analyzed using Volocity software (Perkin Elmer). The plasmids encoding WT mEGFP-HRas (Plasmid #18662) and DN mEGFP-HRas S17N (Plasmid #18665) were purchased from Addgene.

### Live-cell microscopy

CV-1 cells were plated at a density of 1.5 x 10^4^ on Lab-Tek II chambered coverglass slides (Nunc). On the following day, cells were infected with SV40 at MOI of 100. At 20 h.p.i., cells were co-transfected with plasmids encoding Lamp1-RFP and YFP-Rab5 using FuGENE6 (Promega) transfection reagent. Time-lapse microscopy was started 40 h.p.i. Images were acquired every 15 min on a Nikon spinning disk confocal microscope with Nikon perfect focus system and a LiveCell environmental chamber (Pathology Devices). Volocity software (PerkinElmer) and ImageJ were used for 4D image analysis.

### Generation of MKK4 knock-down CV-1 cells with shRNA

Three different shRNAs to MKK4 were generated using the MISSION library (Sigma) and a lentiviral system consisting of pRSV, pMDL and pVSV-G. Briefly, virus was produced by transfection of HEK293 cells with the transfer and packaging vectors using FuGENE6 (Promega). At 24 and 48 h.p.i., virus-containing supernatant was filtered using a 0.45 µm nylon membrane filter and stored at -80°C. Pooled virus preparations were used for CV-1 target cells transduction. About 24 h.p.i., puromycin treatment was started and maintained until the end of experiments. MKK4 knockdown was verified by qPCR.

The following MKK4 shRNAs were used in this study:

A12: TRCN0000001390 (CCGGCTTCTTATGGATTTGGATGTACTCGAGTACATCCAAATCCATAAGAAGTTTTT)

B1: TRCN0000001391

(CCGGGATGTATGAAGAACGTGCCGTCTCGAGACGGCACGTTCTTCATACATCTTTTT)

B3: TRCN0000001393

(CCGGGATATGATGTCCGCTCTGATGCTCGAGCATCAGAGCGGACATCATATCTTTTT)

Scrambled shRNA: SHC002

(CCGGCAACAAGATGAAGAGCACCAACTCGAGTTGGTGCTCTTCATCTTGTTGTTTTT)

### Flow cytometry

Flow cytometry was done as previously described ((37)). Briefly, to stain for intracellular VP1 and large T antigen, cells were fixed and permeabilized with methanol or 4% PFA/0.5% Triton X-100 and subsequently stained with the primary antibodies PAb 597 and PAb108 against VP1 and large T antigen, respectively. Alexa Fluor 488-labeled donkey anti-mouse antibody (Jackson Research) or goat anti-mouse APC were used as secondary antibodies. Data were acquired on an AccuriC6 or FACS Calibur flow cytometer (BD Biosciences) and analyzed with FlowJo software (Treestar).

### LDH release assay

LDH release into the cell culture supernatant was quantified using the CytoTox 96 non-radioactive cytotoxicity assay (Promega) according to the manufacturer’s instructions. Raw data were collected on a spectrophotometer at 490 nm. Values were calculated as follows: OD (infected sample) - OD (mock-infected sample) / OD (infected biological control) - OD (mock-infected biological control).

### Data analysis

All data were primarily processed in Microsoft Excel. Statistical analysis and graph production was performed using Graphpad prism software. For statistical analysis of data, an unpaired t test was used.

## Acknowledgments

We thank Thomas Magaldi for discussions and Jan Zulkeski for assistance in preparing this manuscript. This work was supported by grants from the NIH to D.D. (NS065719 and AI102876).

## FIGURE LEGENDS

**Supplemental Figure S1.**
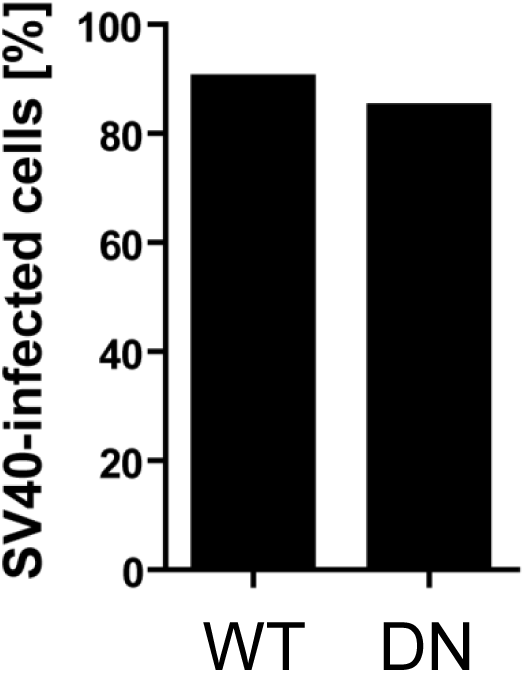
Expression of dominant-negative Ras does not affect infection levels of CV-1 cells. CV-1 cells were transfected with wild-type mEGFP-HRas (WT) or dominant-negative mEGFP-HRas S17N (DN), respectively. Twelve hours p.i., cells were infected with SV40 at MOI of 10. At 24 h.p.i., cells were harvested and stained for large T antigen. Cells were analyzed by flow cytometry for GFP and T antigen fluorescence. The fraction of infected cells in the GFP^+^ population is shown.

**Supplemental Figure S2.**
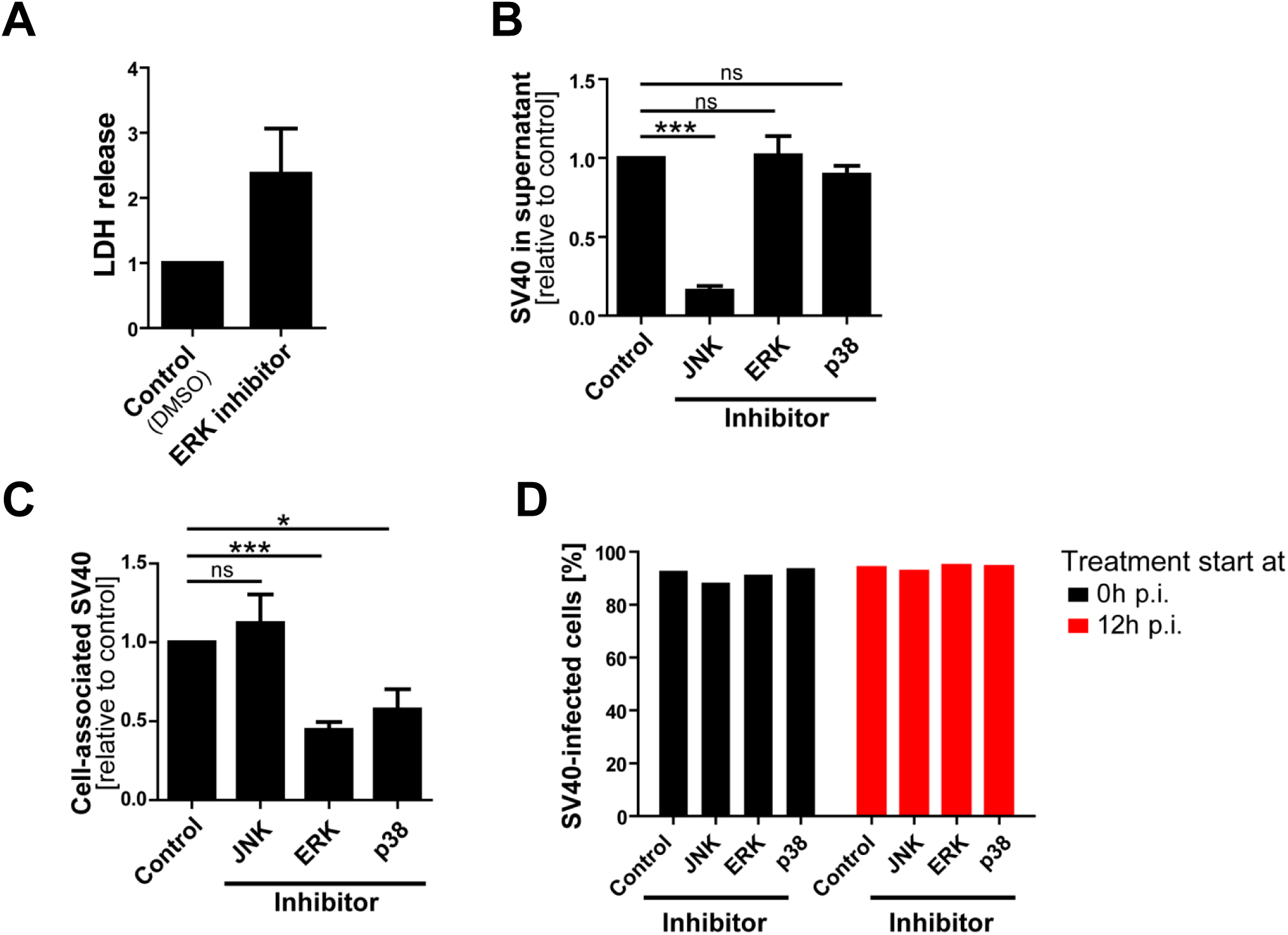
Effect of MAPK inhibitors on SV40 replication and cell viability. (**A)** ERK inhibitor reduces cell viability as assessed by LDH release. Mock-infected CV-1 cells were treated with ERK inhibitor or vehicle alone for 36 h and read out for LDH activity in the cell supernatant. Depicted is the mean of three independent experiments +/- SEM. **(B)** Treatment with JNK Inhibitor SP600125 reduces release of SV40 from infected cells. CV-1 cells were infected at MOI of 10. Inhibitor treatment was started at 12 h.p.i. and infectious units in the supernatant were quantified at 48 h.p.i. (**C)** Treatment with JNK inhibitor does not affect virus production. CV-1 cells were treated as in (**B**). Infectious units were quantified in cells at 48 h.p.i. **(D)** Inhibitor treatment of CV-1 cells does not influence SV40 infectivity. CV-1 cells were infected at MOI of 10 and treated with inhibitors against JNK, ERK, and p38 or DMSO vehicle at the time of infection (0 h.p.i. – black bars) or 12 h.p.i. (red bars). At 24 h.p.i., cells were harvested, stained for expression of large T antigen, and read out by flow cytometry.

**Supplemental Figure S3.**
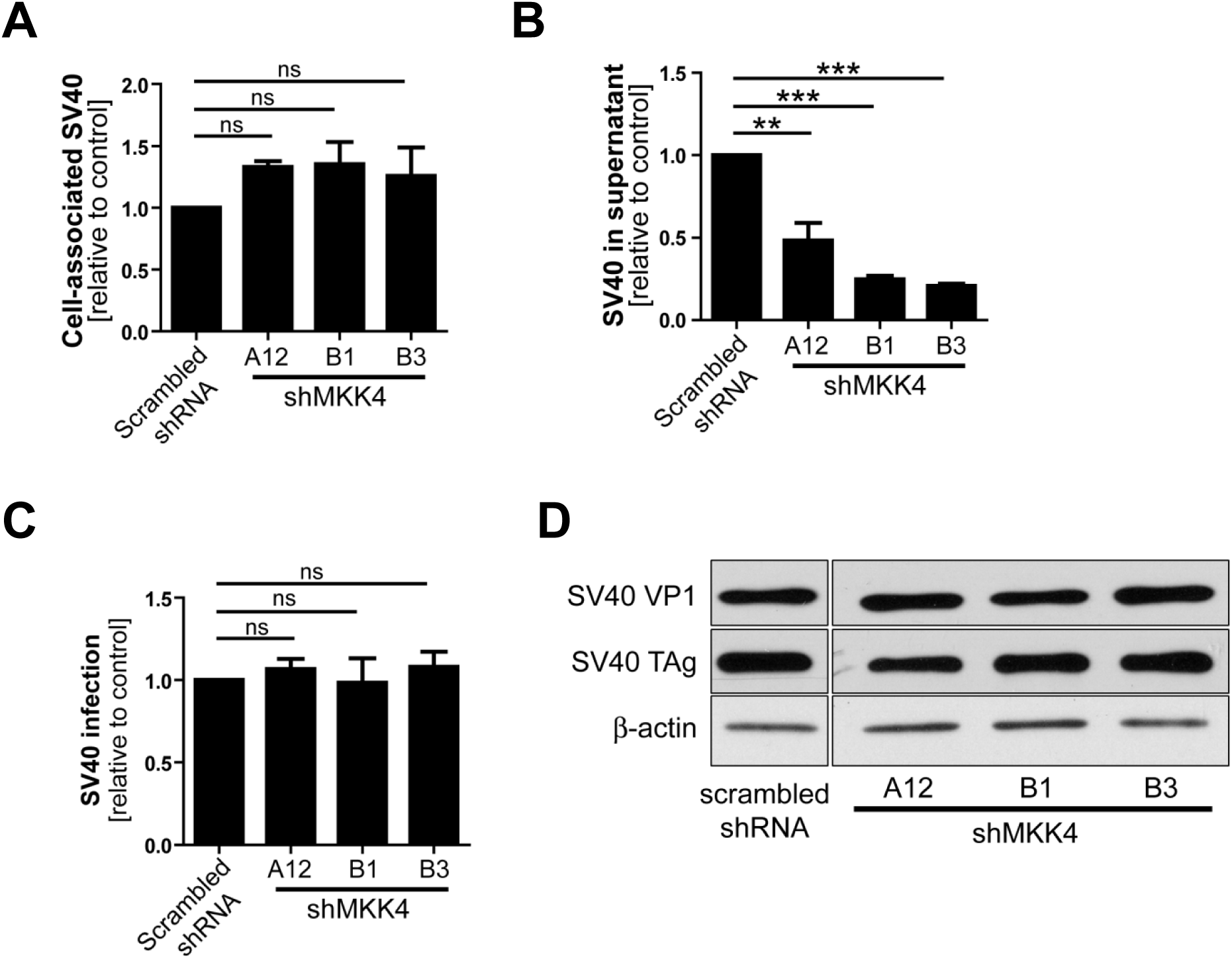
MKK4 knock-down reduces SV40 release while maintaining infection levels and virus replication. **(A)** MKK4 knockdown does not affect virus production. CV-1 cells expressing shRNAs to MKK4 or scrambled shRNA were infected with SV40 at MOI of 10. At 48 h.p.i., cells were harvested for quantitation of infectious units. **(B)** MKK4 is required for efficient virus release. Experiments were set up as in (A). At 48 h.p.i., infectious units were quantified in the supernatant (**C, D)**. MKK4 knockdown does not influence SV40 infectivity or replication. Experiments were set up as described in (**A**). (**C**) At 24 h.p.i. infected cells were harvested and subjected to large T antigen staining for determination of infection levels. Depicted is the mean of three experiments +/-SEM. (**D**) At 48 h.p.i., expression levels of the early protein large T antigen and the late protein VP1 were determined by immunoblotting.

**Supplemental Figure S4.**
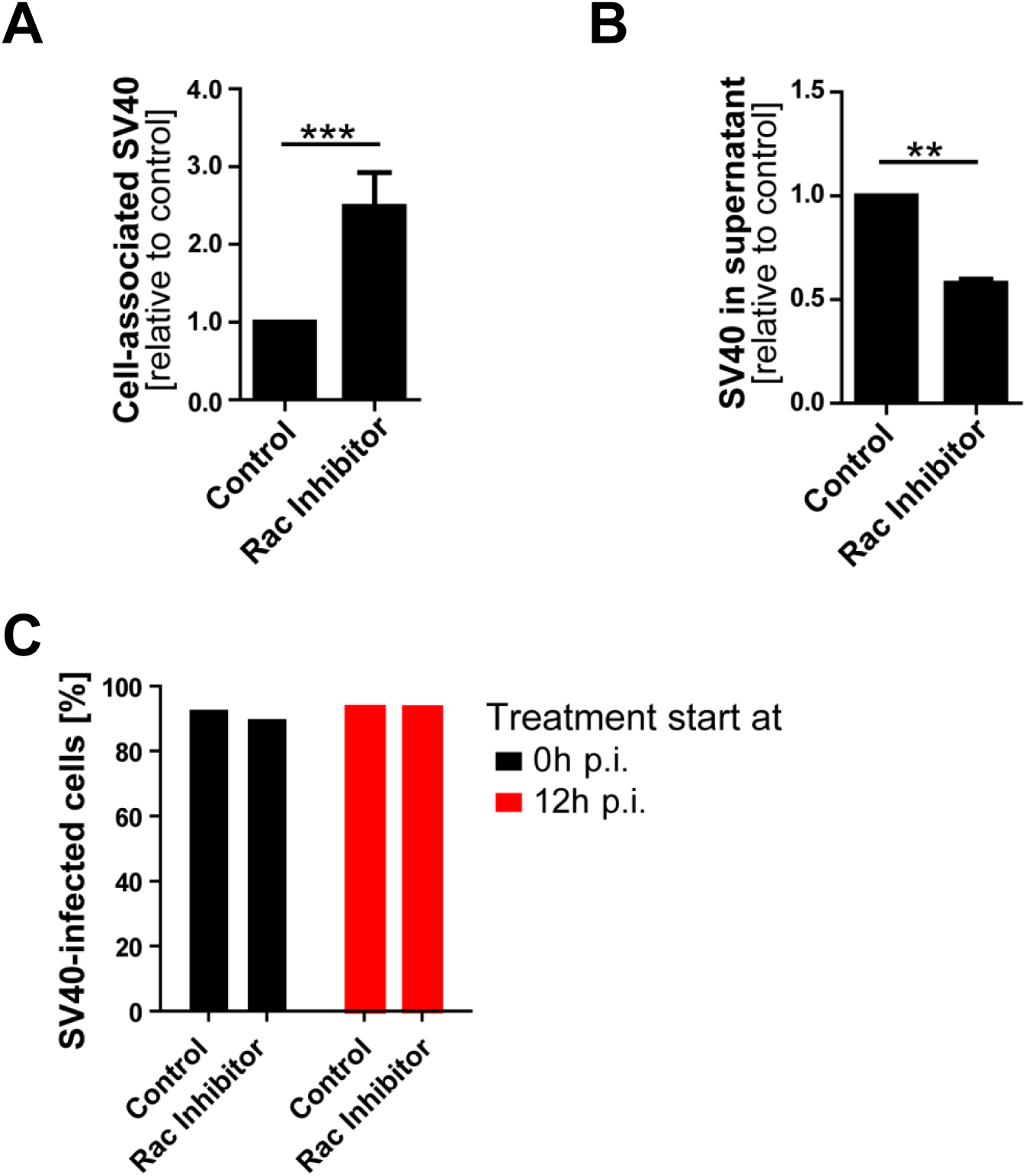
Effect of Rac1 inhibitor EHT1864 on SV40 replication. **(A)** Rac1 inhibition increases levels of cell associated virus. CV-1 cells were infected at MOI of 10. Inhibitor treatment was started at 12 h.p.i. and cell-associated infectious units were quantified at 48 h.p.i. **(B)** Treatment with Rac1 Inhibitor EHT1864 reduces release of SV40 from infected cells. CV-1 cells were treated as in (A). Infectious units in the supernatant were quantified at 48 h.p.i. (**C**) Inhibitor treatment of CV-1 cells does not influence SV40 infectivity. CV-1 cells were infected at MOI of 10 and treated with Rac1 inhibitor EHT1864 or DMSO vehicle at the time of infection (0 h.p.i. – black bars) or 12 h.p.i (red bars). At 24 h.p.i., cells were harvested, stained for expression of large T antigen, and read out by flow cytometry.

## Movie Legends

### Suppl. Movie S1

#### Vacuole Fusion

Time-lapse movie showing fusion of YFP-Rab5-positive vacuoles in an SV40-infected CV1 cell. Rab5-YFP is shown in green; Lamp1-RFP is shown in red.

### Suppl. Movie S2

#### Vacuole dynamics

Time-lapse movie showing dynamics of SV40-induced vacuole maturation in an SV40-infected CV1 cell. Rab5-YFP is shown in green; Lamp1-RFP is shown in red.

## References

1. Dalianis T, Hirsch HH. 2013. Human polyomaviruses in disease and cancer. Virology 437:63-72.

2. Assetta B, Atwood WJ. 2017. The biology of JC polyomavirus. Biol Chem 398:839-855.

3. Kaliyaperumal S, Dang X, Wuethrich C, Knight HL, Pearson C, MacKey J, Mansfield KG, Koralnik IJ, Westmoreland SV. 2013. Frequent infection of neurons by SV40 virus in SIV-infected macaque monkeys with progressive multifocal leukoencephalopathy and meningoencephalitis. Am J Pathol 183:1910-1917.

4. Miskin DP, Koralnik IJ. 2015. Novel syndromes associated with JC virus infection of neurons and meningeal cells: no longer a gray area. Curr Opin Neurol 28:288-294.

5. Dupzyk A, Tsai B. 2016. How Polyomaviruses Exploit the ERAD Machinery to Cause Infection. Viruses 8.

6. Schelhaas M, Malmstrom J, Pelkmans L, Haugstetter J, Ellgaard L, Grunewald K, Helenius A. 2007. Simian Virus 40 depends on ER protein folding and quality control factors for entry into host cells. Cell 131:516-529.

7. Querbes W, Benmerah A, Tosoni D, Di Fiore PP, Atwood WJ. 2004. A JC virus-induced signal is required for infection of glial cells by a clathrin- and eps15-dependent pathway. J Virol 78:250-256.

8. DuShane JK, Wilczek MP, Mayberry CL, Maginnis MS. 2018. ERK Is a Critical Regulator of JC Polyomavirus Infection. J Virol 92.

9. Glenn GM, Eckhart W. 1990. Transcriptional regulation of early-response genes during polyomavirus infection. J Virol 64:2193-2201.

10. O'Hara SD, Garcea RL. 2016. Murine Polyomavirus Cell Surface Receptors Activate Distinct Signaling Pathways Required for Infection. MBio 7.

11. Zullo J, Stiles CD, Garcea RL. 1987. Regulation of c-myc and c-fos mRNA levels by polyomavirus: distinct roles for the capsid protein VP1 and the viral early proteins. Proc Natl Acad Sci U S A 84:1210-1214.

12. Dangoria NS, Breau WC, Anderson HA, Cishek DM, Norkin LC. 1996. Extracellular simian virus 40 induces an ERK/MAP kinase-independent signalling pathway that activates primary response genes and promotes virus entry. J Gen Virol 77 (Pt 9):2173-2182.

13. Pelkmans L, Fava E, Grabner H, Hannus M, Habermann B, Krausz E, Zerial M. 2005. Genome-wide analysis of human kinases in clathrin- and caveolae/raft-mediated endocytosis. Nature 436:78-86.

14. Daniels R, Rusan NM, Wadsworth P, Hebert DN. 2006. SV40 VP2 and VP3 insertion into ER membranes is controlled by the capsid protein VP1: implications for DNA translocation out of the ER. Mol Cell 24:955-966.

15. Henriksen S, Hansen T, Bruun JA, Rinaldo CH. 2016. The Presumed Polyomavirus Viroporin VP4 of Simian Virus 40 or Human BK Polyomavirus Is Not Required for Viral Progeny Release. J Virol 90:10398-10413.

16. Sweet BH, Hilleman MR. 1960. The vacuolating virus, S.V. 40. Proc Soc Exp Biol Med 105:420-427.

17. Luo Y, Motamedi N, Magaldi TG, Gee GV, Atwood WJ, DiMaio D. 2016. Interaction between Simian Virus 40 Major Capsid Protein VP1 and Cell Surface Ganglioside GM1 Triggers Vacuole Formation. MBio 7:e00297.

18. Magaldi TG, Buch MHC, Murata H, Erickson KD, Garcea RL, Peden K, Stehle T, DiMaio D. 2012. Mutations in the GM1 binding site in SV40 VP1 alter receptor usage and cellular tropism. J Virol 86:7028-7042.

19. Miyamura T, Kitahara T. 1975. Early cytoplasmic vacuolization of African green monkey kidney cells by SV40. Arch Virol 48:147-156.

20. Shubin AV, Demidyuk IV, Komissarov AA, Rafieva LM, Kostrov SV. 2016. Cytoplasmic vacuolization in cell death and survival. Oncotarget 7:55863-55889.

21. Nagahama M, Itohayashi Y, Hara H, Higashihara M, Fukatani Y, Takagishi T, Oda M, Kobayashi K, Nakagawa I, Sakurai J. 2011. Cellular vacuolation induced by Clostridium perfringens epsilon-toxin. FEBS J 278:3395-3407.

22. Overmeyer JH, Kaul A, Johnson EE, Maltese WA. 2008. Active ras triggers death in glioblastoma cells through hyperstimulation of macropinocytosis. Mol Cancer Res 6:965-977.

23. Kaul A, Overmeyer JH, Maltese WA. 2007. Activated Ras induces cytoplasmic vacuolation and non-apoptotic death in glioblastoma cells via novel effector pathways. Cell Signal 19:1034-1043.

24. Matallanas D, Arozarena I, Berciano MT, Aaronson DS, Pellicer A, Lafarga M, Crespo P. 2003. Differences on the inhibitory specificities of H-Ras, K-Ras, and N-Ras (N17) dominant negative mutants are related to their membrane microlocalization. J Biol Chem 278:4572-4581.

25. Maltese WA, Overmeyer JH. 2014. Methuosis: nonapoptotic cell death associated with vacuolization of macropinosome and endosome compartments. Am J Pathol 184:1630-1642.

26. Pezzarossa A, Zosel F, Schmidt T. 2015. Visualization of HRas Domains in the Plasma Membrane of Fibroblasts. Biophys J 108:1870-1877.

27. Bhanot H, Young AM, Overmeyer JH, Maltese WA. 2010. Induction of nonapoptotic cell death by activated Ras requires inverse regulation of Rac1 and Arf6. Mol Cancer Res 8:1358-1374.

28. Porat-Shliom N, Kloog Y, Donaldson JG. 2008. A unique platform for H-Ras signaling involving clathrin-independent endocytosis. Mol Biol Cell 19:765-775.

29. Chinnapen DJ, Hsieh WT, te Welscher YM, Saslowsky DE, Kaoutzani L, Brandsma E, D'Auria L, Park H, Wagner JS, Drake KR, Kang M, Benjamin T, Ullman MD, Costello CE, Kenworthy AK, Baumgart T, Massol RH, Lencer WI. 2012. Lipid sorting by ceramide structure from plasma membrane to ER for the cholera toxin receptor ganglioside GM1. Dev Cell 23:573-586.

30. Hancock JF. 2003. Ras proteins: different signals from different locations. Nat Rev Mol Cell Biol 4:373-384.

31. Yamazaki T, Zaal K, Hailey D, Presley J, Lippincott-Schwartz J, Samelson LE. 2002. Role of Grb2 in EGF-stimulated EGFR internalization. J Cell Sci 115:1791-1802.

32. Chiu VK, Bivona T, Hach A, Sajous JB, Silletti J, Wiener H, Johnson RL, 2nd, Cox AD, Philips MR. 2002. Ras signalling on the endoplasmic reticulum and the Golgi. Nat Cell Biol 4:343-350.

33. Clayson ET, Brando LV, Compans RW. 1989. Release of simian virus 40 virions from epithelial cells is polarized and occurs without cell lysis. J Virol 63:2278-2288.

34. Chi S, Kitanaka C, Noguchi K, Mochizuki T, Nagashima Y, Shirouzu M, Fujita H, Yoshida M, Chen W, Asai A, Himeno M, Yokoyama S, Kuchino Y. 1999. Oncogenic Ras triggers cell suicide through the activation of a caspase-independent cell death program in human cancer cells. Oncogene 18:2281-2290.

35. Raghava S, Giorda KM, Romano FB, Heuck AP, Hebert DN. 2011. The SV40 late protein VP4 is a viroporin that forms pores to disrupt membranes for viral release. PLoS Pathog 7:e1002116.

36. Stroh LJ, Maginnis MS, Blaum BS, Nelson CD, Neu U, Gee GV, O'Hara BA, Motamedi N, DiMaio D, Atwood WJ, Stehle T. 2015. The Greater Affinity of JC Polyomavirus Capsid for alpha2,6-Linked Lactoseries Tetrasaccharide c than for Other Sialylated Glycans Is a Major Determinant of Infectivity. J Virol 89:6364-6375.

37. Goodwin EC, Lipovsky A, Inoue T, Magaldi TG, Edwards APB, Van Goor KEY, Paton AW, Paton JC, Atwood WJ, Tsai B, DiMaio D. 2011. BiP and multiple DNAJ molecular chaperones in the endoplasmic reticulum are required for efficient simian virus 40 infection. mBio 2:101-111.

